# BETA- AND GAMMA-SYNUCLEINS MODULATE SYNAPTIC VESICLE-BINDING OF ALPHA-SYNUCLEIN

**DOI:** 10.1101/2020.11.19.390419

**Authors:** Kathryn E. Carnazza, Lauren Komer, André Pineda, Yoonmi Na, Trudy Ramlall, Vladimir L. Buchman, David Eliezer, Manu Sharma, Jacqueline Burré

## Abstract

α-Synuclein (αSyn), β-synuclein (βSyn), and γ-synuclein (γSyn) are abundantly expressed in the vertebrate nervous system. αSyn functions in neurotransmitter release via binding to and clustering synaptic vesicles and chaperoning of SNARE-complex assembly. The functions of βSyn and γSyn are unknown. Functional redundancy of the three synucleins and mutual compensation when one synuclein is deleted have been proposed, but with conflicting evidence. Here, we demonstrate that βSyn and γSyn have a reduced affinity towards membranes compared to αSyn, and that direct interaction of βSyn or γSyn with αSyn results in reduced membrane binding of αSyn. Our data suggest that all three synucleins affect synapse function, but only αSyn mediates the downstream function of vesicle clustering and SNARE-complex assembly, while βSyn and γSyn modulate the activity of αSyn through regulating its binding to synaptic vesicles.

## INTRODUCTION

α-Synuclein (αSyn), β-synuclein (βSyn) and γ-synuclein (γSyn) are abundantly expressed proteins in the vertebrate nervous system (Buchman et al., 1998b; George, 2002; Jakes et al., 1994; Ji et al., 1997; Lavedan et al., 1998; Nakajo et al., 1993). αSyn plays an important physiological role at the synapse, to maintain neurotransmitter release via binding to synaptic vesicles (Iwai et al., 1995; Kahle et al., 2000; Maroteaux et al., 1988), regulating synaptic vesicle pools (Cabin et al., 2002; Murphy et al., 2000; Yavich et al., 2004), and by chaperoning of SNARE-complex assembly (Burré et al., 2010). Aggregation of αSyn is a pathological hallmark of Parkinson’s disease, Lewy body dementia, multiple system atrophy, and a variety of other synucleinopathies (Spillantini et al., 1997). Despite the involvement of βSyn and γSyn in some neurodegenerative diseases including Lewy body dementia, diffuse Lewy body disease, Gaucher’s disease, and Parkinson’s disease (Galvin et al., 2000; Galvin et al., 1999; Nguyen et al., 2011; Ninkina et al., 2009; Nishioka et al., 2010; Peters et al., 2012; Surgucheva et al., 2002), virtually nothing is known about their physiological functions in the brain.

The three synucleins share a high degree of sequence homology (Jakes et al., 1994; Nakajo et al., 1993), are highly expressed in the human brain and show a strikingly similar regional distribution. Synucleins are coexpressed in the cerebral cortex, amygdala, caudate nucleus, substantia nigra, striatum, hippocampus, and thalamus (Ahmad et al., 2007; Jeannotte et al., 2009; Lavedan, 1998; Lavedan et al., 1998; Murphy et al., 2000; Ninkina et al., 1998), while in other brain areas, their expression patterns differ. For example, the subthalamic nucleus expresses mainly γSyn whereas the corpus callosum has αSyn and γSyn, but no βSyn (Lavedan, 1998). In regions where co-expression occurs, the relative levels of synucleins differ. Evolutionarily recent regions express predominantly αSyn and βSyn, whereas more ancient structures have higher levels of γSyn expression (Buchman et al., 1998b; Jakes et al., 1994; Jakowec et al., 2001; Lavedan, 1998; Maroteaux and Scheller, 1991; Ueda et al., 1993; Ueda et al., 1994). αSyn is most abundant in the frontal cortex, hippocampus, olfactory bulb, thalamus, and striatum (Iwai et al., 1995), while less αSyn is found in the more caudal regions, such as brainstem and spinal cord. In contrast, βSyn is predominantly expressed in the neocortex, hippocampus, striatum, thalamus, and cerebellum (George, 2002). In the brain, highest expression of γSyn is found in the substantia nigra, hippocampus, thalamus, caudate nucleus, amygdala, and particularly, the motor nuclei of the brain stem (Buchman et al., 1998b; Ji et al., 1997; Lavedan et al., 1998).

These expression patterns suggest that each synuclein may partake in a distinct function, as suggested by some studies, although functional redundancy has been proposed as well. A compensatory function is supported by increased expression of the remaining family member in the CNS of α/β-Syn and α/γ-Syn double knockout mice (Chandra et al., 2004; Robertson et al., 2004), and accelerated pathology in CSPα knockout mice in the absence of both αSyn and βSyn compared to αSyn only (Chandra et al., 2005). Functional redundancy is supported by a behavioral and dopamine release phenotypes in α/γ-Syn knockout mice while single αSyn KO or γSyn KO mice showed no deficits (Senior et al., 2008), by considerable functional overlap in the pools of genes whose expression is changed in the absence of αSyn or γSyn (Kuhn et al., 2007), and by a decrease in dopamine levels in brains of α/β-Syn KO but not αSyn or βSyn single KO (Chandra et al., 2004).

In contrast, other studies found no compensatory increase in αSyn or βSyn expression upon knockout of γSyn (Ninkina et al., 2003; Papachroni et al., 2005), in βSyn upon knockout of αSyn and γSyn (Papachroni et al., 2005), or in βSyn or γSyn upon αSyn knockout (Abeliovich et al., 2000; Kuhn et al., 2007; Schluter et al., 2003). Lack of compensation was also surmised from lack of exaggeration of phenotypes in α/γ-Syn double KO mice compared to αSyn and γSyn single KO mice (Robertson et al., 2004). Redundancy was countered by lack of potentiation of gene expression in α/γ-Syn knockout versus αSyn and γSyn single knockout mice and differentially regulated gene expression in αSyn and γSyn KO mice (Kuhn et al., 2007), and a reduction in striatal dopamine and specific protein levels in αSyn null but not γSyn null mice (Al-Wandi et al., 2010), a lack of γSyn ability to interact with VAMP2, support SNARE-complex assembly and rescue the CSPα knockout phenotype (Ninkina et al., 2012), as well as a distinct effect of the lack of only βSyn in an inverted grid test, and a distinct effect of the lack of only αSyn on striatal dopamine levels that was not exacerbated by additional loss of the other synucleins (Connor-Robson et al., 2016). These discrepancies may be either due to the fact that different brain regions and different animal ages were analyzed in these studies, or they may point to a different role of the three synucleins in the same cellular process.

Here, we demonstrate the first important physiological function for βSyn and γSyn that can explain and consolidate the seemingly controversial findings above. Our data support a model where all synuclein family members affect synapse function, but their specific roles in that process differ, in that only αSyn mediates the downstream function of vesicle clustering and SNARE-complex assembly while βSyn and γSyn modulate the activity of αSyn through regulating its binding to synaptic vesicles.

## RESULTS

### βSyn and γSyn reveal reduced α-helical content and binding to liposomes compared to αSyn

αSyn binds to synaptic vesicle membranes (Iwai et al., 1995; Kahle et al., 2000; Maroteaux et al., 1988), reflecting its preference for membranes with high curvature (Davidson et al., 1998; Middleton and Rhoades, 2010). βSyn and γSyn share the highly conserved α-helical lipid-binding motif with αSyn (Figure S1A), and can adopt the same two-helix conformation (Rivers et al., 2008; Sung and Eliezer, 2006), suggesting that they bind to lipids as well. Yet, the lipid binding domain of βSyn shares only 87% sequence identity with αSyn and lacks 11 residues toward the end, and γSyn shares only 68% sequence identity in the lipid binding domain (Figure S1A). These differences may lead to different membrane binding properties.

When we analyzed the ability of αSyn, βSyn, or γSyn to associate with artificial liposomes of 30 nm diameter in a flotation assay (Figure S1B), we found robust binding of αSyn to liposomes composed of 30% phosphatidylserine and 70% phosphatidylcholine, as well as to liposomes mimicking synaptic vesicles (Figure 1A; Figure S1C). Strikingly, βSyn and γSyn revealed a dramatically reduced binding to both of these liposomes, which was not affected by deletion of the C-termini that are not important for the interaction with membranes (Figure 1A; Figure S1C-E).

**Figure 1.**
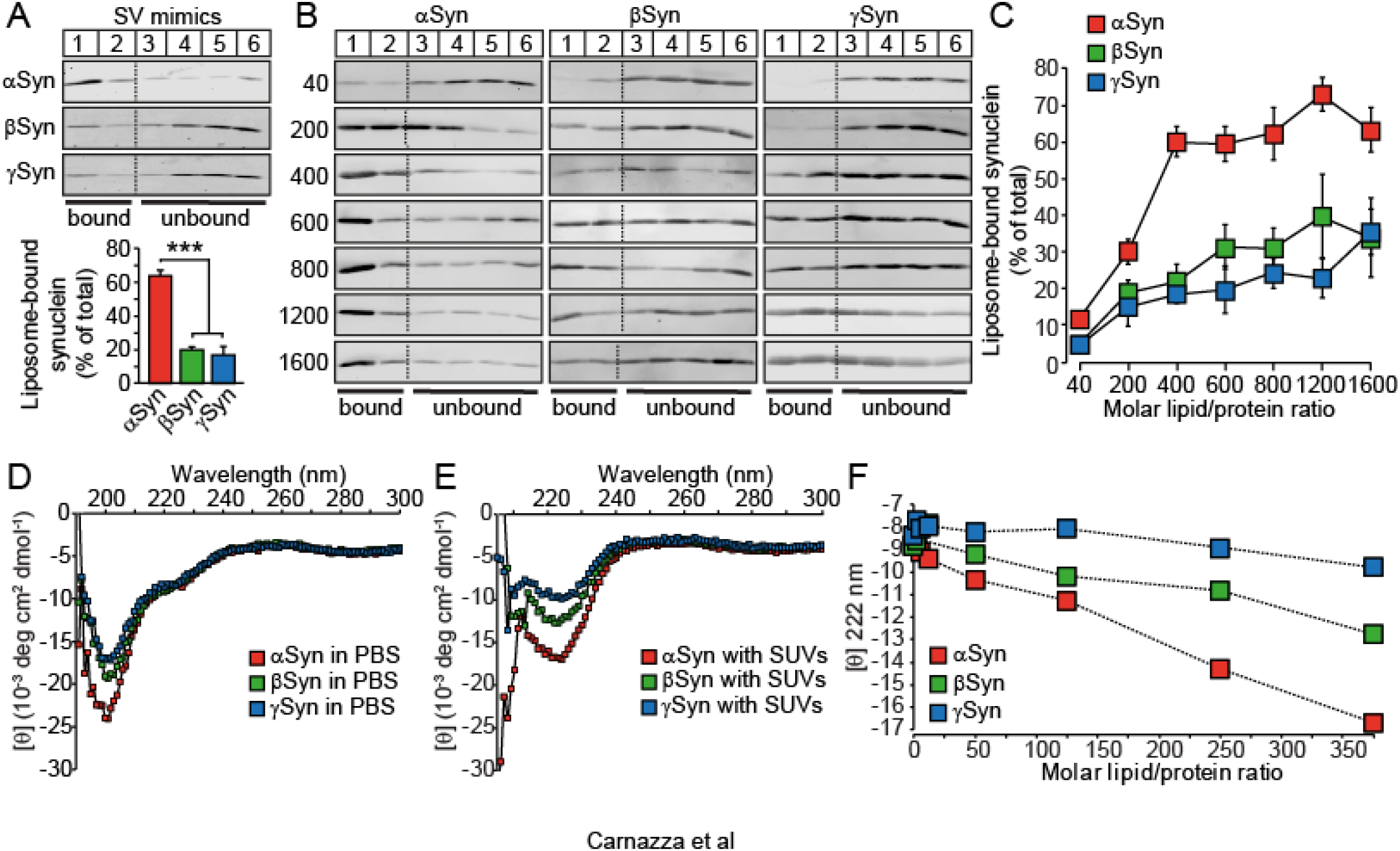
Binding of recombinant synucleins to liposomes. (A-C) αSyn, βSyn or γSyn were floated by density gradient centrifugation with synaptic vesicle mimics (30 nm diameter; composition: 36% PC, 30% PE, 12% PS, 5% PI, 7% SM, 10% cholesterol) or artificial small unilamellar vesicles (30 nm diameter; composition: 70% PC, 30% PS). Top two fractions 1 and 2 were defined as lipid-bound fractions (see Figure S1B). Flotation of synucleins with liposomes was quantified as the sum of the top 2 fractions, plotted as the percentage of total synuclein in the gradient. (B,C) Same as in (A), except that different molar lipid/protein ratios of small unilamellar vesicles were used. Data are means ± SEM (***p < 0.001 by Student’s t test; n = 6 - 8 independent experiments). (D-F) CD spectroscopy of synucleins. (D, E) Secondary structure of recombinant αSyn, βSyn or γSyn in absence (D) or presence (E) of 30 nm SUVs at a molar lipid/protein ratio of 400. (F) Same as (E), except that different molar/lipid ratios were used, and signal at 222 nm was plotted to highlight α-helicity (mean of n = 3).

αSyn preferentially associates with liposomes of high curvature (Davidson et al., 1998; Middleton and Rhoades, 2010), but βSyn and γSyn may have a preference for less curved membranes. We thus repeated our flotation assay in presence of liposomes of 100 nm (Figure S1D) or 200 nm (Figure S1E) diameter, but lost binding of all synucleins to these larger liposomes.

Last, αSyn, βSyn, and γSyn may have different affinities and thus potentially different saturation levels towards liposomes. We thus tested binding of synucleins to liposomes at different molar lipid:protein ratios, extending beyond the known saturation levels for αSyn (Figure 1B and 1C). We found saturation of binding of all synuclein family members around a molar lipid:protein ratio of 400. Strikingly, βSyn and γSyn binding plateaued at a significantly lower liposome-bound protein percentage compared to αSyn (Figure 1B and 1C).

αSyn is natively unstructured in solution, but adopts an α-helical conformation upon binding to membranes (Burre et al., 2013; Davidson et al., 1998; Iwai et al., 1995; Kahle et al., 2000; Maroteaux et al., 1988). Thus, as a separate means to assess binding of synucleins to liposomes, we used circular dichroism (CD) spectroscopy. While all synucleins had a similar unfolded nature in solution (Figure 1D), βSyn and γSyn revealed reduced α-helicity at all molar lipid:protein ratios tested (Figure 1E and 1F). These data are in agreement with our liposome flotation data (Figure 1A-C; Figure S1) and suggest that βSyn and γSyn have significantly reduced affinities towards membranes compared to αSyn.

These data suggest a reduced ability of βSyn and γSyn to associate with phospholipid membranes, which is in agreement with the literature reporting a reduced propensity of βSyn for α-helical secondary structure in the N-terminal region compared to αSyn (Bertoncini et al., 2007; Sung and Eliezer, 2006), a five-fold lower affinity of βSyn binding to DMPS vesicles relative to αSyn (Brown et al., 2016), a two-fold lower affinity for βSyn and γSyn binding to 1:1 POPC:POPS vesicles relate to αSyn (Ducas and Rhoades, 2012; Middleton and Rhoades, 2010), and a reduced binding affinity of βSyn to POPA:POPC vesicles compared to αSyn (Sharma et al., 2020).

### Reduced presynaptic localization of γSyn but not βSyn compared to αSyn

The flotation and CD experiments above used purified recombinant synucleins and phospho-liposomes. To probe for membrane binding in a more physiological context, we performed subcellular fractionation on wild-type (WT) mouse brains to separate brain homogenate into cytosolic and membrane-bound fractions. We found the integral membrane protein VDAC1 and membrane-associated protein SNAP-25 to be robustly associated with membrane fractions, demonstrating successful fractionation (Figure S2A). Synucleins were more cytosolic, representative of their on/off membrane equilibrium. Compared to αSyn, we found a significant reduction in membrane-bound βSyn and γSyn, with αSyn > γSyn > βSyn detected in the bound fraction (Figure S2).

αSyn targets to presynaptic terminals via binding to synaptic vesicle membranes (Iwai et al., 1995; Kahle et al., 2000; Maroteaux et al., 1988) and to the synaptic vesicle protein synaptobrevin-2/VAMP2 (Burre et al., 2012; Burré et al., 2010). Subcellular compartment-specific membrane and/or protein interactions may thus affect the synaptic localization of all synucleins. We thus enriched for synaptosomes from brain homogenates of WT mice and analyzed relative synuclein levels in synaptosomes. We found an enrichment for the synaptic vesicle proteins synaptobrevin-2 and CSPα, while α-tubulin and neurofilament of 165 kDa were depleted, as expected (Figure 2A and 2B). When we analyzed the synucleins, we found neither enrichment nor depletion of αSyn and βSyn in synaptosomes compared to brain homogenates, while γSyn showed a significant depletion in synaptosomes compared to total brain homogenate (Figure 2A and 2B). In parallel, we analyzed the synaptic colocalization of αSyn, βSyn or γSyn with the synaptic marker synapsin in primary cortical WT mouse neurons. We found a robust synaptic localization for αSyn and βSyn, similar to the integral synaptic vesicle proteins SV2 and synaptobrevin-2, while γSyn was not as synaptic (Figure 2C-E). These findings agree with the reported presynaptic localization for αSyn and βSyn, and the more widely dispersed localization for γSyn throughout the cytosol (Buchman et al., 1998a; Buchman et al., 1998b; Jakes et al., 1994; Maroteaux et al., 1988; Mori et al., 2002; Murphy et al., 2000; Surguchov et al., 2001).

**Figure 2.**
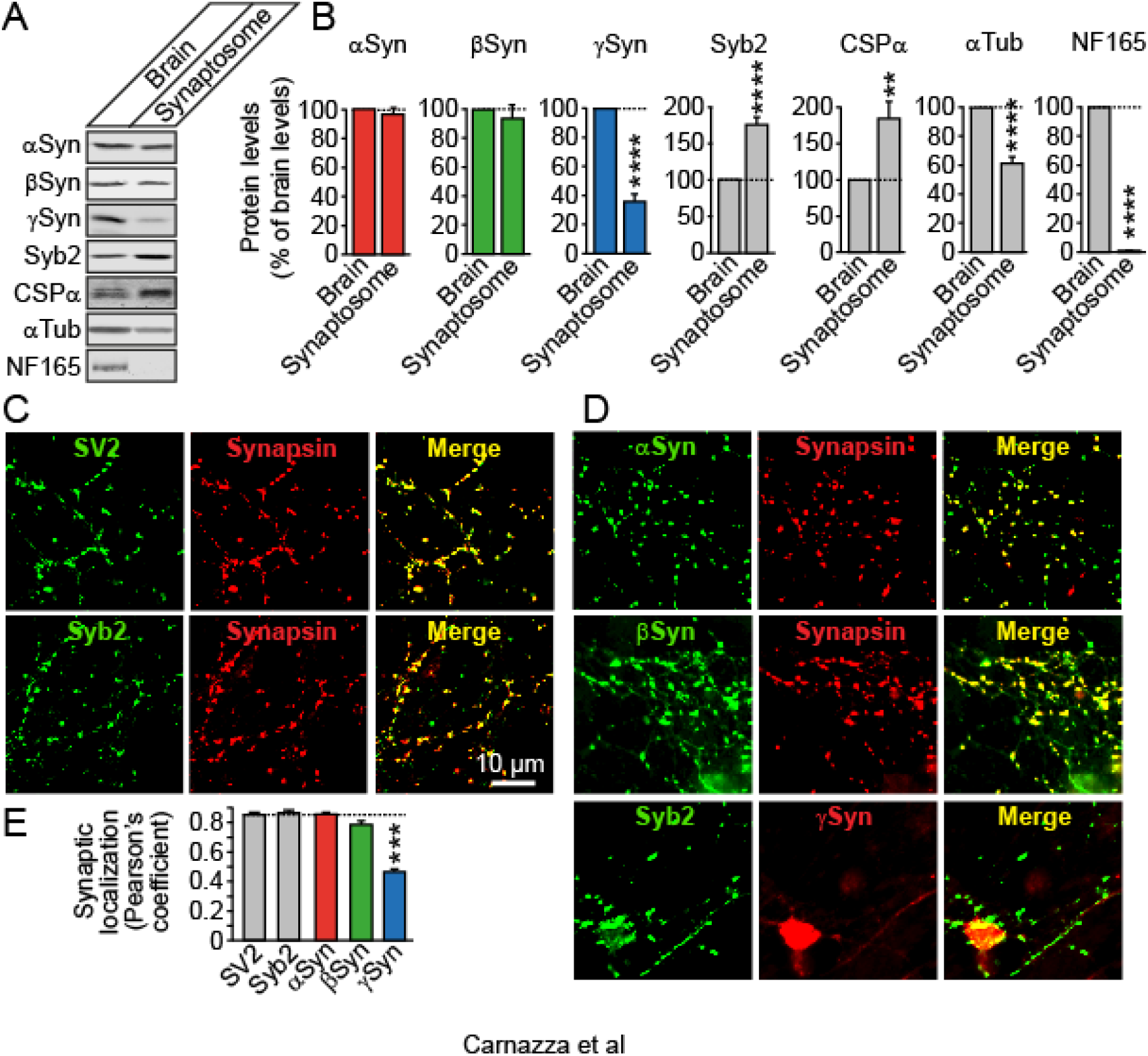
Presynaptic localization of synucleins. (A, B) Brains of P40 WT mice were homogenized and half of the homogenate was subjected to subcellular fractionation to yield synaptosomes. Equal protein amount of brain homogenate and synaptosomes was separated by SDS-PAGE and analyzed by quantitative immunoblotting of the indicated protein (Syb2, synaptobrevin-2; αTub, α-tubulin; NF165, neurofilament of 165 kDa). (C-E) Primary cortical WT mouse neurons were analyzed at 27DIV for the indicated proteins. Colocalization was quantitated using Pearson’s coefficient. Data are means ± SEM ((** p < 0.01, *** p < 0.001 by Student’s t test; n = 4 brains in (A, B) and 6 independent cultures in (C-E)).

### Synucleins interact with each other in a specific conformation

Despite a dramatically reduced ability to associate with membranes compared to αSyn, how do βSyn and, to a lesser extent, γSyn, still target to the synapse? Based on our previous studies showing homo-multimerization of the membrane-binding domain of αSyn (Burre et al., 2014), synucleins may interact with each other, thereby enabling synaptic co-localization. In fact, there is precedence for a possible *in vivo* interaction among the synuclein family members in the literature, although mainly focused on inhibition of αSyn aggregation by βSyn or γSyn (Biere et al., 2000; Hashimoto et al., 2001; Park and Lansbury, 2003; Uversky et al., 2002; Windisch et al., 2002): (1) βSyn affects the speed of the nucleation phase of αSyn during in vitro fibrillation (Brown et al., 2016; Van de Vondel et al., 2018), (2) αSyn/βSyn heterodimers were found in yeast (Tenreiro et al., 2016), (3) NMR PRE titration experiments measured a Kd for the αSyn/βSyn heterodimer of 100 μM, while the αSyn/αSyn homodimer Kd’s were 500 μM (Janowska et al., 2015), (4) molecular dynamics and potential of mean force computational study found that it is more favorable for αSyn to complex with βSyn to form heterodimers, rather than a second αSyn to form homodimers (Sanjeev and Mattaparthi, 2017; Tsigelny et al., 2007), and (6) weak to moderate μM binding affinities were found for αSyn-βSyn, βSyn-γSyn, and αSyn-γSyn, with the αSyn-βSyn interaction being the weakest (Jain et al., 2018).

When we probed directly for interactions among the synuclein family members using co-immunoprecipitation in absence of membranes, we did not detect any binding (Figure S3A). Given that multimers of αSyn only form upon membrane binding of αSyn (Burre et al., 2014), αSyn-βSyn and αSyn-γSyn interactions may require membranes as well. Yet, immunoprecipitation in presence of membranes is complicated by non-specific binding and thus not feasible. We thus adapted our FRET system that we had previously used to assess the specific configuration of αSyn multimers on membranes (Burre et al., 2014). For this, we introduced cysteine residues at the beginning and the end of the α-helical domains of the synucleins, generated recombinant proteins, and labeled these cysteines either with Alexa488 or Alexa546 (Figure 3A). Based on our previous studies (Burre et al., 2014), we assumed that not only homo-multimers but also hetero-multimers of synucleins would adopt the antiparallel broken helix configuration (Fig. 3B), and thus labeled only residues within the synucleins that match this configuration (Fig. 3A, B). We then measured FRET between various combinations in presence or absence of liposomes (Figure 3C, D; Figure S3). Only in presence of charged liposomes did we detect a FRET signal between αSyn, βSyn or γSyn, suggesting that all synucleins share the ability to form homo-dimers (Figure 3C). As expected, we were unable to detect any FRET signal between synucleins in presence of neutral liposomes that synucleins do not bind to. Importantly, our FRET experiments also provide evidence for hetero-multimerization between the synucleins in the presence of membranes (Figure 3D).

**Figure 3.**
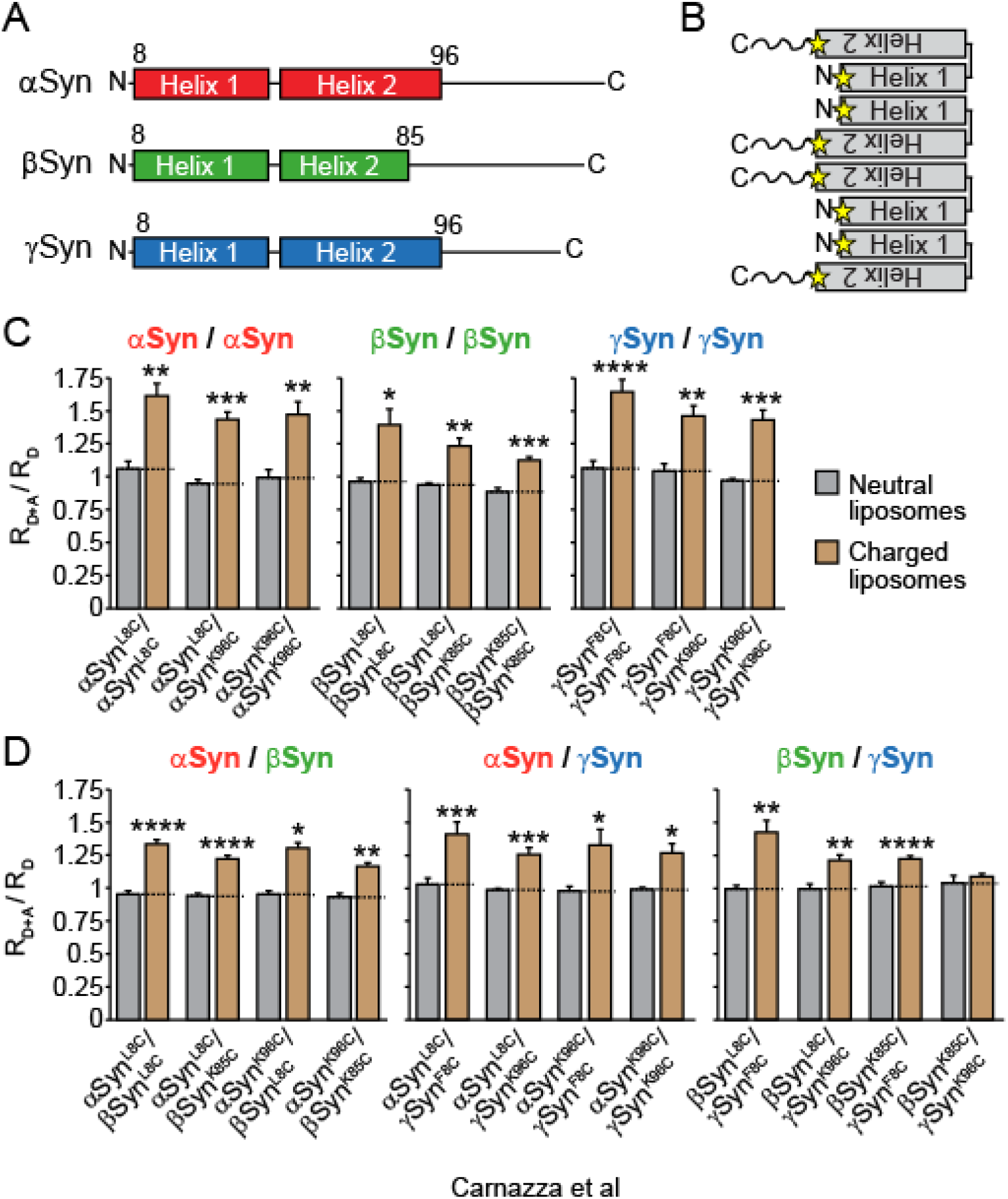
Interaction between synucleins. (A) Fluorescent synuclein labeling scheme for FRET experiments. Single-cysteine substitutions were introduced into synucleins at position 8 and 96 for αSyn and γSyn, and at position 8 and 85 for βSyn for modification with Alexa 488- or Alexa546-maleimide. (B) Configuration of synuclein multimers (stars = labeled residues). (C, D) Emission spectra in Figure S3C-H were used for calculation of FRET signals between synucleins. Data are means ± SEM (* p < 0.05, ** p < 0.01, *** p < 0.001 by Student t test; n = 3 independent experiments).

### Synucleins modulate each other’s ability to associate with membranes

Does hetero-multimerization of synucleins have a functional consequence? For physiological implications of this interaction, synucleins need to be expressed in the same neuron. When we tested if αSyn is co-expressed with βSyn or γSyn in wild-type neurons, we found βSyn to highly co-localize with αSyn in presynaptic compartments (indicated by individual puncta), while the overlap of γSyn with αSyn was largely restricted to dendritic and somatic compartments (Figure S4A and S4B). Note that not all presynaptic terminals stain equally strong for each of the synucleins, indicating heterogeneity in synucleins levels in different neurons.

We then analyzed if presence or absence of βSyn or γSyn results in changes in the synaptic targeting of αSyn, using co-expression via lentiviral vectors in αβγ-synuclein triple knockout neurons. When we quantified synaptic targeting of αSyn with increasing levels of βSyn or γSyn, we found a significant dose-dependent reduction in synaptic localization of αSyn (Figure 4A-D; Figure S4C-F).

**Figure 4.**
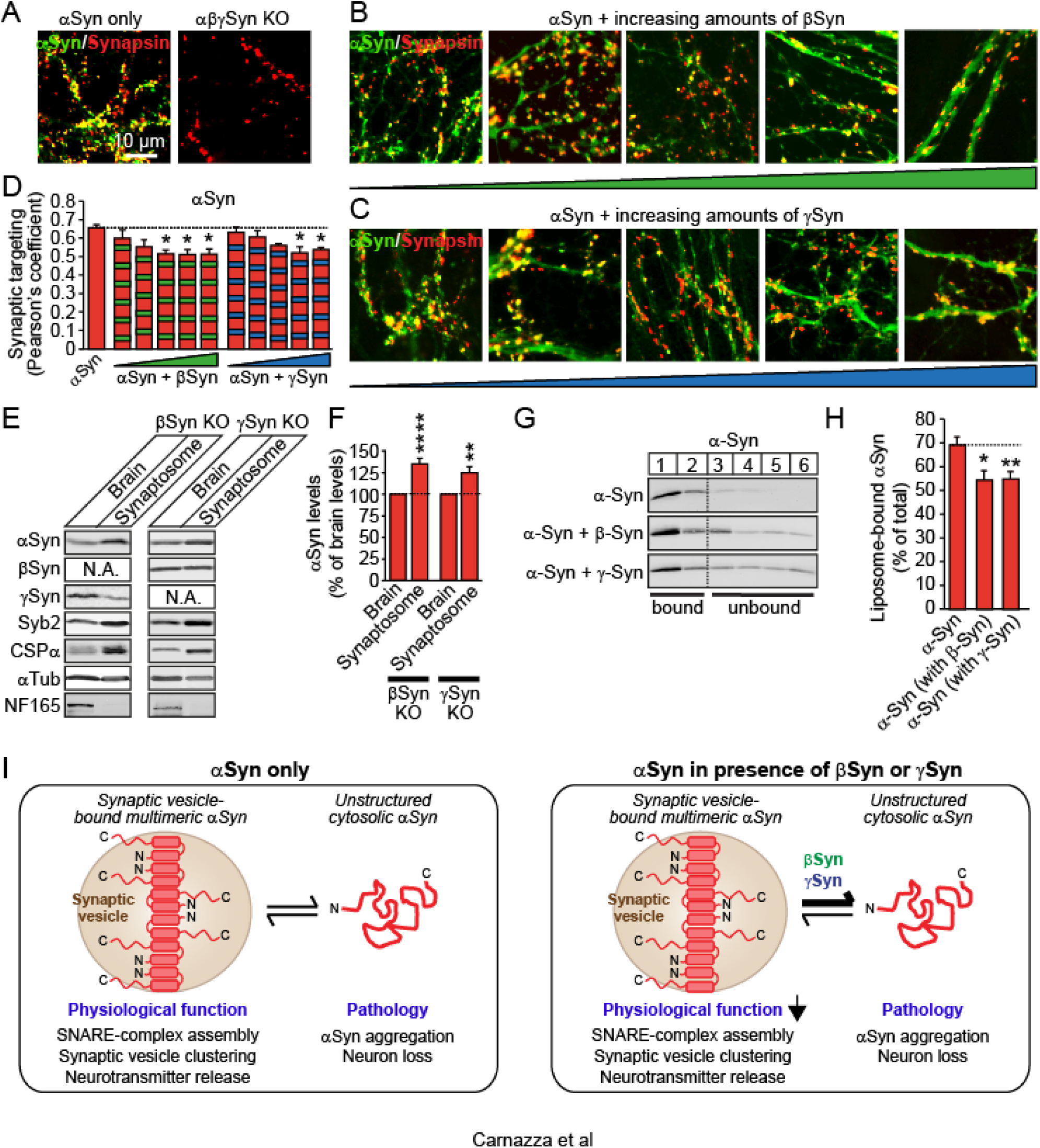
Decreased synaptic vesicle binding of αSyn in presence of βSyn or γSyn. (A-D) Synaptic targeting of αSyn in absence or presence of increasing amounts of βSyn or γSyn. αβγ-Syn triple knockout neurons were transduced with lentiviral vectors expressing αSyn only (A) or αSyn with increasing amounts of βSyn (B) or γSyn (C). Synaptic targeting was quantified by co-localization with synapsin and Pearson’s coefficient (D). Data are means ± SEM (* p < 0.05 by Student t test; n = 3 cultures). (E, F) Analysis of αSyn enrichment in synaptosomes. Synaptosomes were isolated from mouse brain homogenates of mice lacking βSyn or γSyn via subcellular fractionation. 20 μg protein of homogenate and synaptosomes were analyzed by quantitative immunoblotting to the indicated proteins (Syb2, synaptobrevin-2; αTub, α-tubulin; NF165, neurofilament of 165 kDa). Data are means ± SEM (** p < 0.01, **** p < 0.0001 by Student t test; n = 6-8 mice; see also Figure S5C-E). (G, H) Co-flotation of synucleins. Liposome binding of αSyn was analyzed in absence or presence of equal amounts of βSyn or γSyn by a flotation assay. Flotation of αSyn with liposomes was quantified as the sum of the top 2 fractions, plotted as the percentage of total synuclein in the gradient. Data are means ± SEM (* p < 0.05, ** p < 0.01 by Student’s t test; n = 7-15 independent experiments). (I) Proposed model of the effect of βSyn or γSyn on αSyn function. Via hetero-multimerization, βSyn and γSyn reduce synaptic vesicle-bound αSyn which leads to a reduction in αSyn’s physiological activity to mediate SNARE-complex assembly, synaptic vesicle clustering and neurotransmitter release.

In parallel, to test synaptic targeting at endogenous expression levels, we generated single αSyn, βSyn, and γSyn knockout mice from our αβγ-synuclein triple knockout mice (Figure S5A and S5B). We then repeated the analysis of synaptic targeting of αSyn, βSyn, and γSyn using a synaptosome enrichment study. When we compared the relative amount of synucleins in synaptosomes versus total brain, we found a significantly increased amount of synaptic αSyn in absence of βSyn or γSyn (Figure 4E and 4F), unlike in WT brains (Figure 2A). Importantly, this was neither due to changes in efficiency of subcellular fractionation, as marker proteins revealed the same enrichment and depletion as in wild-type brains (Figure 2A and 2B, and Figure S5C.E), nor was it due to altered expression levels of synucleins in these genotypes (Figure S5A and S5B).

To test directly if βSyn and γSyn reduce the ability of αSyn to associate with synaptic vesicle membranes, we measured liposome binding of αSyn in absence or presence of equal amounts of βSyn or γSyn and vice versa using a liposome flotation assay. Interestingly, we found a significant reduction in membrane association of αSyn in presence of βSyn or γSyn (Figure 4G and 4H), and an increase in membrane association of βSyn and γSyn when αSyn was present (Figure S5F and S5G), suggesting that synucleins directly affect each other’s ability to bind to synaptic vesicle membranes.

Overall, our data suggest that via hetero-multimerization, the largely cytosolic βSyn and γSyn reduce the amount of synaptic vesicle-bound αSyn, potentially providing a tuning mechanism for αSyn function in synaptic vesicle clustering and neurotransmitter release (Figure 4H).

## DISCUSSION

The function of αSyn is tightly linked to its localization in presynaptic terminals. Targeting of αSyn to terminals is mediated by binding to synaptic vesicle lipids and synaptobrevin-2 (Burré et al., 2010; Iwai et al., 1995; Kahle et al., 2000; Maroteaux et al., 1988; Sun et al., 2019), and binding to vesicles triggers multimerization of αSyn which in concert with binding of αSyn to synaptobrevin-2, promotes synaptic vesicle clustering (Diao et al., 2013; Sun et al., 2019). The implications of this clustering activity have been suggested to restrict synaptic vesicle mobility between synapses (Scott and Roy, 2012) and to serve as a reserve pool of synaptic vesicles for long-term operation of a neuron during high-frequency stimulation (Diao et al., 2013). In agreement, loss of synucleins increases tethering of synaptic vesicles to the active zone and reduces links between vesicles (Vargas et al., 2017), while interlocked αSyn/synaptobrevin-2 dimers reduce dispersion (Sun et al., 2019). In addition, Ca^2+^ has been suggested to regulate the interaction of αSyn with synaptic vesicles (Lautenschlager et al., 2018), and electron microscopy studies show a redistribution of αSyn with activity (Tao-Cheng, 2006).

Here, we have identified a new control mechanism for synaptic vesicle-binding of αSyn (Figure 4H). The physiological functions of βSyn and γSyn have remained largely elusive, although a neuroprotective role for both has been proposed (da Costa et al., 2003; Hashimoto et al., 2001; Windisch et al., 2002), suggesting that interactions among synucleins are important and need to be examined. We demonstrate here that βSyn and γSyn have a direct effect on the physiologically functional pool of αSyn on synaptic vesicles, suggesting that the roles of synucleins are complementary, but not functionally redundant, which may explain some of the controversial findings in the field. How is this process regulated within a presynaptic terminal where two or three synucleins co-localize? The decrease of αSyn interaction with the synaptic vesicle membrane could be due to direct competition of the synucleins for binding sites on the synaptic vesicle membrane. In this case, more αSyn would bind to the vesicle surface in neurons lacking βSyn or γSyn due to lack of competition. However, βSyn and γSyn reveal a reduced binding affinity towards membranes (Figures 1 and 2), so unless there is an excess amount of βSyn or γSyn in a terminal, there may not be significant competition. In addition, presence of αSyn increases membrane binding of βSyn and γSyn (Figure S5), which also argues against competition. As an alternative mechanism, binding of βSyn or γSyn to αSyn on the synaptic vesicle surface may reduce the affinity of αSyn for synaptic vesicle membranes, either by interfering with the αSyn-membrane interaction directly, or by reducing the overall binding affinity of αSyn for membranes through the formation of heteromers. Regardless of the specific mechanism, our findings raise several important questions that are essential for our understanding of synuclein biology and pathology and will need to be addressed in follow-up studies:

The synucleins are co-expressed at varied levels in the brain (Buchman et al., 1998b; George, 2002; Iwai et al., 1995; Jakes et al., 1994; Jakowec et al., 2001; Ji et al., 1997; Lavedan, 1998; Lavedan et al., 1998; Maroteaux and Scheller, 1991; Ueda et al., 1993; Ueda et al., 1994), and single cell RNA seq analysis of mouse cortex and hippocampus reveals expression ratios of αSyn/βSyn between 0.64-2.3, with the majority between 1-1.5, and αSyn/γSyn ratios between 1.1-1.85, with a lack of detection of γSyn in most cells (Allen Institute for Brain Science)(Hawrylycz et al., 2012; Lein et al., 2007). In agreement, analyses of human cortex reveals expression ratios of αSyn/βSyn between 0.002-6.31, with the majority between 1-1.5, and αSyn/γSyn ratios between 0.088-362, also with a lack of detection of γSyn in most cells (Allen Institute for Brain Science) (Hawrylycz et al., 2012; Lein et al., 2007). This vast heterogeneity suggests that relative ratios of αSyn, βSyn, and γSyn could modulate synaptic vesicle clustering and SNARE-complex assembly and thus neuron function via regulating the relative amount of αSyn on synaptic vesicles. αSyn has been shown to disperse from the nerve terminal in response to neural activity (Fortin et al., 2005). Furthermore, expression of αSyn is high in regions of the adult CNS that display synaptic plasticity (Maroteaux and Scheller, 1991), and in adult canaries and zebra finches, where αSyn expression correlates with plasticity in the developing song control system (Clayton and George, 1999). At present, it remains to be determined whether plastic regions in the brain feature altered expression levels of αSyn, βSyn, and γSyn.

Besides identifying a new physiological function for βSyn and γSyn, our findings may also have implications for the contributions of βSyn and γSyn to disease. βSyn and γSyn have links to Lewy body dementia, diffuse Lewy body disease, Gaucher’s disease, motor neuron disease, neurodegeneration with brain iron accumulation type 1, glaucoma, and Parkinson’s disease (Galvin et al., 2000; Galvin et al., 1999; Nguyen et al., 2011; Ninkina et al., 2009; Nishioka et al., 2010; Peters et al., 2012; Surgucheva et al., 2002). We and others have previously shown that lack of SNARE chaperoning by αSyn or the CSPα chaperone complex causes progressive neuropathology and premature death (Burre et al., 2012, 2015; Burré et al., 2010; Greten-Harrison et al., 2010; Sharma et al., 2012; Sharma et al., 2011), and synuclein triple knockout mice exhibit an age- and activity-dependent decrease in SNARE-complex assembly which correlates with progressive neuropathology and leads to premature death (Burré et al., 2010). Thus, αSyn is required for the long-term sustenance of nerve terminals and neurons and protects against age-dependent decline by ensuring SNARE-complex assembly over the long life of a neuron. Rendering αSyn less membrane-associated through the presence of βSyn and γSyn may increase the aggregation-prone cytosolic pool of αSyn. Alternatively, binding of αSyn to βSyn or to γSyn may shield the aggregation-prone residues in αSyn. The relative ratios of the three synucleins may explain the selective vulnerability of certain neuronal populations to dysfunction and degeneration.

Finally, our findings may point to an alternative therapeutic strategy by adjusting not only αSyn levels, but focusing on βSyn and γSyn levels as well. Previous strategies have concentrated largely on eliminating αSyn from the brain. Yet, removal of αSyn can cause functional changes to neuronal synapses in various *in vivo* experimental systems, particularly in the aged nervous system (Al-Wandi et al., 2010; Benskey et al., 2018; Collier et al., 2016; Gorbatyuk et al., 2010; Markopoulou et al., 2014; Ninkina et al., 2020; Robertson et al., 2004), and strategies targeting βSyn or γSyn to alter aggregation-prone pools of αSyn may be less detrimental.

In summary, our data suggest that a correct balance of synucleins is important for normal brain function, and that an imbalance of these proteins might not only affect neuron function and plasticity, but also neuronal survival.

## METHODS

### Mouse strains

WT and synuclein null mice were maintained on a C57BL/6 background. Synuclein triple knockout mice were maintained as described previously (Burré et al., 2010). αSyn, βSyn and γSyn single knockout mouse lines were generated by crossing the synuclein triple knockout mice to wild type C57BL/6 mice, and then back crossing the triple-hemizygous progeny to wild type progeny for 5 generations, before separating each of the synuclein knockout alleles. Mice of either sex were used for primary neuronal culture, and no inclusion criteria were used. Mice were housed with a 12-h light/dark cycle in a temperature-controlled room with free access to water and food. All animal procedures were performed according to NIH guidelines and approved by the Committee on Animal Care at Weill Cornell Medicine.

### Cell culture and maintenance

HEK293T cells (ATCC) were maintained in DMEM with 1% penicillin and streptomycin and 10% bovine serum. For production of lentiviral vectors, cells were transfected with equimolar amounts of lentiviral vector FUW containing myc-tagged or untagged αSyn, βSyn or γSyn, pMD2-G-VSVg, pMDLg/pRRE, and pRSV-Rev using calcium phosphate produced in house. 1 h prior to transfection, 25 μM chloroquine in fresh media was added. DNA was incubated for 1 min at room temperature in 100 mM CaCl_2_ and 1× HBS (25 mM HEPES pH 7.05, 140 mM NaCl, and 0.75 mM Na_2_HPO_4_) and the transfection mix was then slowly added to the cells. Medium was replaced with fresh medium after 6 h. Medium containing the viral particles was collected 48 h later and centrifuged for 10 min at 500 g_av_ to remove cellular debris. Viral particles were subsequently concentrated tenfold by centrifugation. Mouse cortical neurons were cultured from newborn mice of either sex. Cortices were dissected in ice-cold HBSS, dissociated and triturated with a siliconized pipette, and plated onto 6 mm poly l-lysine-coated coverslips (for immunofluorescence) or on 24-well plastic dishes. Plating media (MEM supplemented with 5 g/l glucose, 0.2 g/l NaHCO_3_, 0.1 g/l transferrin, 0.25 g/l insulin, 0.3 g/l l-glutamine, and 10% fetal bovine serum) was replaced with growth media (MEM containing 5 g/l glucose, 0.2 g/l NaHCO_3_, 0.1 g/l transferrin, 0.3 g/l l-glutamine, 5% fetal bovine serum, 2% B-27 supplement, and 2 μM cytosine arabinoside) 2 days after plating. At 6 days in vitro (DIV), neurons were transduced with recombinant lentiviruses expressing synucleins. Neurons were harvested or used for experiments as indicated at 27 DIV.

### Immunoprecipitation

Transfected HEK293T cells were solubilized in PBS, pH 7.4, containing 0.15% Triton X-100 and protease inhibitors (Roche). Following centrifugation at 16,000 g_av_ for 20 min at 4°C, the clarified lysate was used for immunoblotting (after addition of 2x SDS sample buffer containing 100 mM DTT) or subjected to immunoprecipitation. Immunoprecipitation was performed with the indicated primary antibodies and 50 μl of a 50% slurry of protein-A sepharose beads (GE Healthcare) for 2 h at 4°C. Control immunoprecipitations were performed with preimmune sera. Following five washes with 1 ml of the extraction buffer, bound proteins were eluted with 2x SDS sample buffer containing 100 mM DTT and boiled for 20 min at 100°C. Coprecipitated proteins were separated by SDS-PAGE, with 5% of the input in the indicated lane.

### Subcellular fractionation

For cytosol/membrane fractionations, entire mouse brains were homogenized in PBS containing protease inhibitors. The homogenates were centrifuged for 1 h at 300,000 g_av_. The supernatant was collected and an equal volume of PBS was added to the pellet. Same volumes were analyzed via SDS-PAGE. Synaptosomes were isolated as previously described (Burre et al., 2006). Briefly, entire mouse brains were homogenized in preparation buffer (5 mM Tris-HCl, 320 mM sucrose, pH 7.4), supplemented with protease inhibitors. The homogenate was centrifuged for 10 min at 1,000 g_av_. The supernatant was collected and the pellet was resuspended in preparation buffer and recentrifuged. Both supernatants were pooled and the final pellet was discarded. Discontinuous Percoll gradients were prepared by layering 7.5 ml supernatant onto three layers of 7.5 ml Percoll solution (3%, 10%, and 23% v/v in 320 mM sucrose, 5 mM Tris-HCl, pH 7.4). After centrifugation for 7 min at 31,400 g_av_, fractions containing synaptosomes were collected, diluted in four volumes of preparation buffer and centrifuged for 35 min at 20,000 g_av_.

### SDS-PAGE and quantitative immunoblotting

Protein samples were separated by SDS-PAGE and either stained using Coomassie Brilliant Blue, or transferred onto nitrocellulose membranes. Blots were blocked in Tris-buffered saline (TBS) containing 0.1% Tween-20 (TBS-T) containing 5% fat-free milk for 30 min at room temperature. The blocked membrane was incubated overnight in PBS containing 1% BSA and 0.2% NaN_3_ and the primary antibody. The blots were then washed twice in TBS-T containing 5% fat-free milk, then incubated for 1 h in the same buffer containing secondary antibody at room temperature. Blots were then washed 3 times in TBS-T, twice in water, and then dried in the dark. Blots were imaged using a LI-COR Odyssey CLx, and images were analyzed using ImageStudioLite (LI-COR).

### Antibodies

CSPα (R807, gift from Dr. Thomas C. Südhof), GAPDH (G-9, Santa Cruz), myc (9E10, deposited to the DSHB by Bishop, J.M.), VDAC1 (N152B/23, Neuromab), NF-165 (2H3, deposited to the DSHB by Jessell, T.M. / Dodd, J.), SNAP-25 (SMI81, Sternberger Monoclonals), synapsin (E028, gift from Dr. Thomas C. Südhof), synaptobrevin-2 (69.1, Synaptic Systems), αSyn (clone 42, BD Biosystems), βSyn (sc-136452, Santa Cruz), γSyn (SK23 (Ninkina et al., 2003)), and α-Tubulin (12G10, DSHB).

### Expression vectors

Full-length human αSyn, βSyn or γSyn cDNA was inserted into modified pGEX-KG vectors (GE Healthcare), containing an N-terminal TEV protease recognition site, or lentiviral vector FUW, without or with an N-terminal myc-tag and a four amino acid linker, resulting in the following N-terminal sequence (EQKLISEEDL-GSGS). Mutant αSyn, βSyn, or γSyn constructs were generated by site-specific mutagenesis, according to the protocol of the manufacturer (Stratagene).

### Recombinant protein expression and purification

All proteins were expressed as GST fusion proteins in bacteria (BL21 strain), essentially as described (Burré et al., 2010). Bacteria were grown to OD 0.6 (measured at 600 nm), and protein expression was induced with 0.05 mM isopropyl β-o-thiogalactoside for 6 h at room temperature. Bacteria were harvested by centrifugation for 20 min at 2,100 g, and pellets were resuspended in solubilization buffer [PBS, 0.5 mg/mL lysozyme, 1 mM PMSF, DNase, and an EDTA-free protease inhibitor mixture (Roche)]. Cells were broken by sonication, and insoluble material was removed by centrifugation for 30min at 7,000 g_av_ and 4 °C. Proteins were affinity-purified using glutathione Sepharose bead (GE Healthcare) incubation overnight at 4 °C, followed by TEV protease (Invitrogen) cleavage overnight at room temperature. His-tagged TEV protease was removed by incubation with Ni-NTA (Qiagen) overnight at 4 °C. Protein concentrations were assessed using the bicinchoninic acid method according to the manufacturer’s protocol (Thermo Scientific).

### Liposome preparation and liposome binding assay

Liposomes were prepared as previously described (Burré et al., 2010). For lipid-binding assays, a mixture of lipids (all Avanti Polar Lipids) in chloroform were dried in a glass vial under a nitrogen stream. Residual chloroform was removed by lyophilization for 2 h. Small unilamellar vesicles were formed by sonicating in PBS on ice. For lipid binding studies, synucleins were incubated with liposomes for 2 h at room temperature at a molar lipid/protein ratio of 400 or other ratios where indicated. For co-flotation experiments, αSyn, βSyn and γSyn were added at a molar lipid/protein ratio of 200 each. Samples were then subjected to a liposome flotation assay (Burré et al., 2010).

### FRET experiments

100 μM GST-fusion protein of αSyn, βSyn or γSyn containing a cysteine (positions 8 and 96 for αSyn and γSyn, positions 8 and 85 for γSyn) were captured on GS4B beads (GE Healthcare). GST-synucleins were reduced with 1 mM DTT for 20 min at 4 °C. Beads were washed four times with PBS containing protease inhibitors and protein were labeled with 2 mM Alexa 488 C5 maleimide or Alexa 546 C5 maleimide (Invitrogen) overnight at 4 °C in the dark. Beads were washed four times with PBS to remove residual unbound dye, and synuclein was eluted from the GST moiety using TEV protease overnight at room temperature. His-tagged TEV protease was removed using Ni-NTA agarose (Qiagen). For FRET experiments, 2.5 μg of Alexa 488-labeled donor synuclein and 2.5 μg of Alexa 546-labeled acceptor synuclein were incubated with and without 100 μg of liposomes in 100 μl of PBS for 2 h at room temperature in the dark. Emission spectra were measured using a Synergy H1 plate reader (BioTek; excitation: 490 nm; emission: 500-650 nm). FRET signals were measured by calculating ratios of RD+A (i.e. ratio of fluorescence of donor plus acceptor α-synuclein at 573 nm and of fluorescence of donor plus acceptor α-synuclein at 519 nm, each subtracted by fluorescence of acceptor α-synuclein) and RD (i.e., ratio of fluorescence of donor α-synuclein at 573 nm and of fluorescence of donor α-synuclein at 519 nm).

### Circular dichroism spectroscopy

Circular dichroism (CD) spectra were measured on an AVIV 62 DS spectrometer equipped with a sample temperature controller. Far-UV CD spectra were monitored from 190 to 300 nm using final protein concentrations of 50 μM and 0.1 mM to 15 mM small unilamellar vesicles (composition: 70% phosphatidylcholine, 30% phosphatidylserine; diameter: ~30 nm) with a path length of 0.2 mm, response time of 1 s, and scan speed of 50 nm/min. Each scan was repeated three times.

### Immunocytochemistry

Cells were washed twice with phosphate-buffered saline (PBS) containing 1 mM MgCl_2_ and were fixed with 4% paraformaldehyde in PBS for 20 min at room temperature. Cells were washed twice with PBS and permeabilized with 0.1% Triton X-100 in PBS for 5 min at RT. After washing twice with PBS, cells were blocked for 20 min with 5% bovine serum albumin (BSA) in PBS. Primary antibody was added in 1% BSA in PBS over night at 4 °C. The next day, cells were washed twice in PBS, blocked for 20 min in 5% BSA in PBS and incubated with secondary antibody and DAPI in 1% BSA in PBS for 1 h at RT in the dark. Cells were washed twice with PBS and were mounted using Fluoromount-G. Cells were imaged on an Eclipse 80i upright fluorescence microscope (Nikon).

### Quantification and statistical analysis

Sample sizes were chosen based on preliminary experiments or similar studies performed in the past. For quantification of immunoblots, a minimum of three independent experiments were performed. For quantification of immunofluorescence microscopy images, images were recorded under the same microscope settings (objective lens and illumination intensity) to ensure reliable quantification across samples and images. Merged images were created using Photoshop (Adobe), and were analyzed using ImageJ (NIH) or Image Studio (LI-COR). No samples or animals were excluded from the analysis, and quantifications were performed blindly. All data are presented as the mean ±SEM, and represent a minimum of three independent experiments. Statistical parameters, including statistical analysis, significance, and n value are reported in each figure legend. Statistical analyses were performed using Prism 8 Software (GraphPad). For statistical comparison of two groups, either two-tailed Student’s t test or two-way ANOVA followed by Bonferroni post hoc test was performed, as indicated in the figure legends. Based on previous studies, a normal distribution of data was assumed, with similar variance between groups that were compared. A value of p < 0.05 was considered statistically significant.

## ACKNOWLEDGEMENTS

We thank Dr. Thomas C. Südhof for providing antibodies. This work was supported by the Russian Science Foundation (Grant 19-14-00064 to V.L.B.), the Alzheimer’s Association (NIRG-15-363678 to M.S.), AFAR (M.S.), the NIH (R37-AG019391 and R35-GM136686 to D.E.; 1R01-AG052505 and 1R01-NS095988 to M.S.; R01-NS102181 and R01-NS113960 to J.B.), and the Leon Levy Foundation (J.B.).

## AUTHOR CONTRIBUTIONS

K.E.C, L.K., A.P., Y.N., M.S., and J.B. designed the study, performed experiments, and analyzed the data, except for the CD studies which were designed and performed by T.R. and D.E.. V.B. provided the γSyn antibody. J.B. wrote the manuscript with input from all authors. All authors discussed and commented on the final manuscript.

