## Supplementary Information for "BETA- AND GAMMA-SYNUCLEINS MODULATE SYNAPTIC VESICLE-BINDING OF ALPHA-SYNUCLEIN"

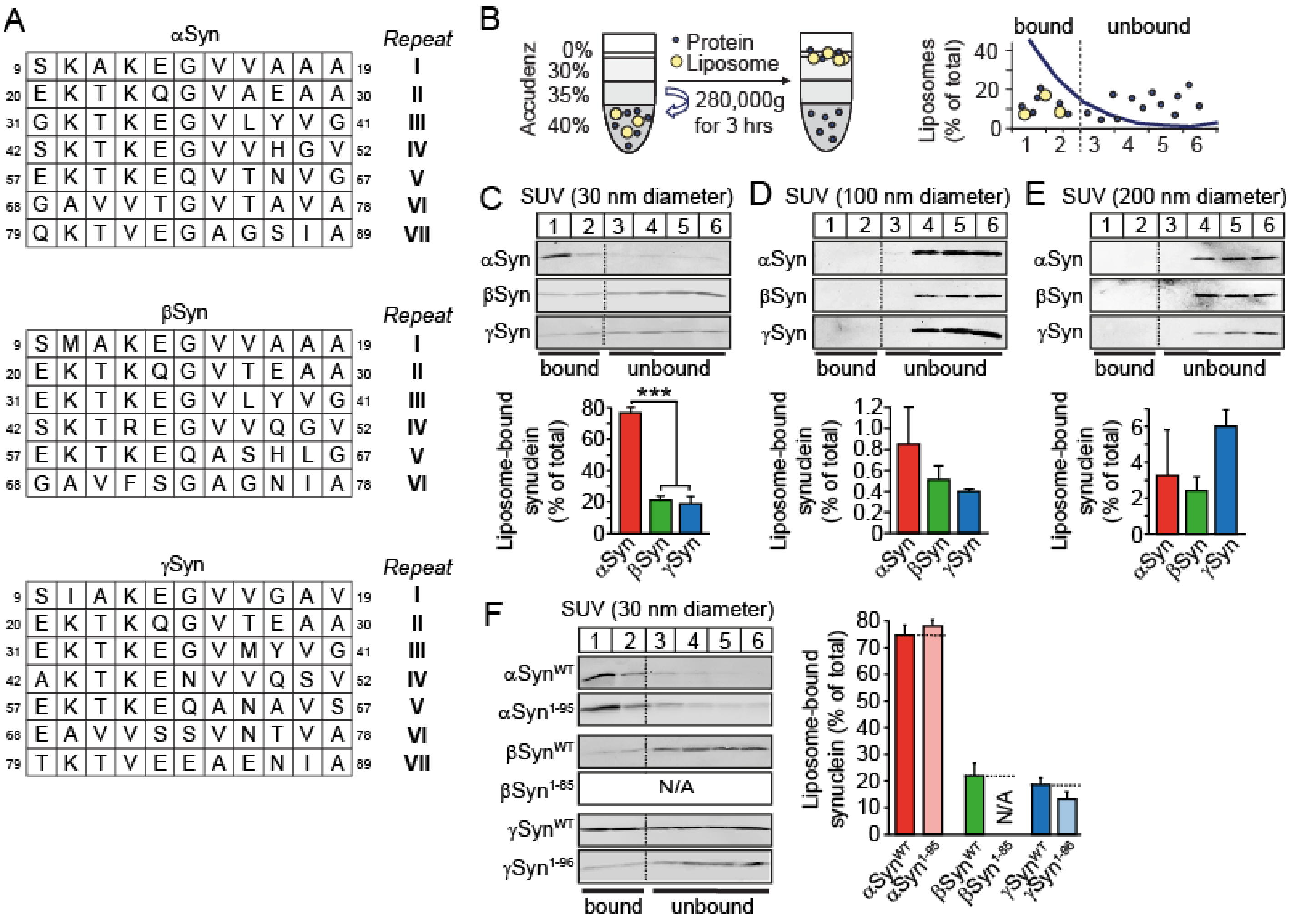


**Supplementary Figure S1. Binding of recombinant synucleins to liposomes.**

(A) Sequence analysis of synucleins. Sequence alignment of αSyn, βSyn, and γSyn to indicate the 11mer repeats that mediate binding of synucleins to membranes.

(B) Phospholipid binding assay. Liposomes mixed with synuclein were floated by density gradient centrifugation. Based on the liposome distribution in the gradient, the top two fractions 1 and 2 were defined as lipid-bound fractions.

(C-F) αSyn, βSyn or γSyn were floated by density gradient centrifugation with small unilamellar vesicles (composition: 70% PC, 30% PS) of 30 nm (C, F), 100 nm (D), or 200 nm (E). Flotation of synucleins with liposomes was quantified as the sum of the top 2 fractions, plotted as the percentage of total synuclein in the gradient. N/A = not analyzed because this truncation mutant did not express. Data are means ± SEM (***$p < 0.001$ by Student's t test; n = 3 - 4 independent experiments).

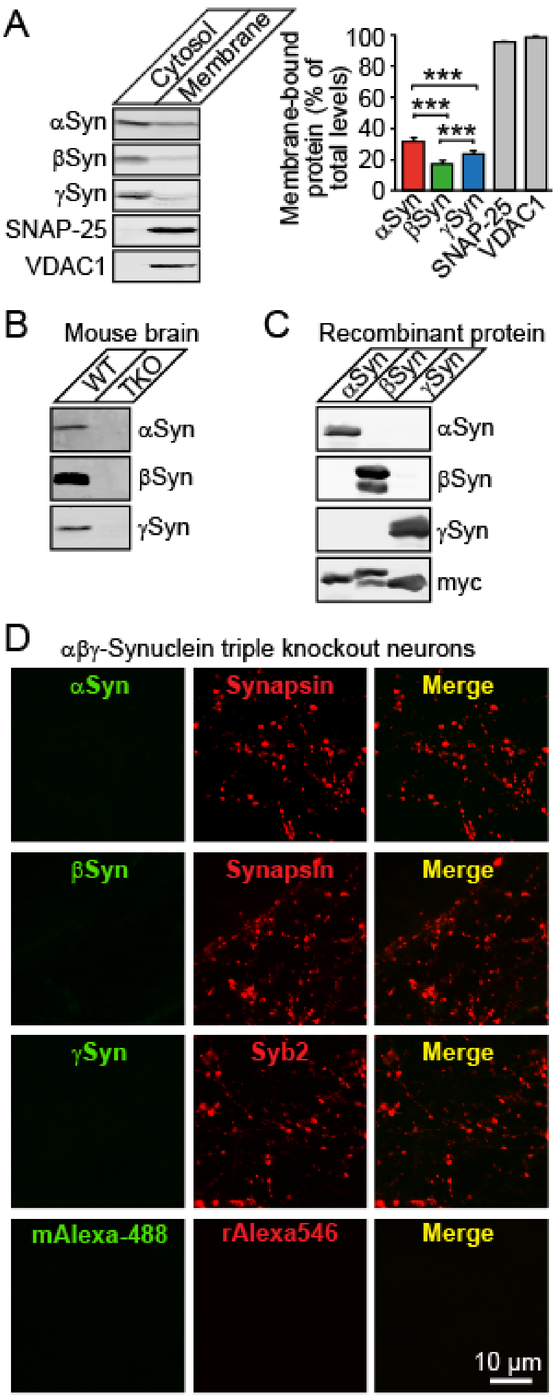


**Supplementary Figure S2. Membrane binding of synucleins.**
(A) Membrane binding of synucleins. P30 WT brains were homogenized and subjected to subcellular fractionation to separate proteins into cytosolic and membrane fractions. Equal amount of protein was separated by SDS-PAGE and subjected to quantitative immunoblotting to the indicated protein. Data are means ± SEM (*** $p < 0.001$ by Student's t test; n = 4 brains).
(B-D) Antibody specificity. 30 µg of brain homogenate of P30 WT and αβγ-synuclein triple knockout mice (B) or 5 µg of myc-tagged recombinant αSyn, βSyn, or γSyn (C) were separated by SDS-PAGE and analyzed by quantitative immunoblotting of the indicated protein (αSyn, BD Bioscience; βSyn, Santa Cruz; γSyn, SK23). Separately, same antibodies were used to stain primary neurons generated from αβγ-synuclein triple knockout mice (D).

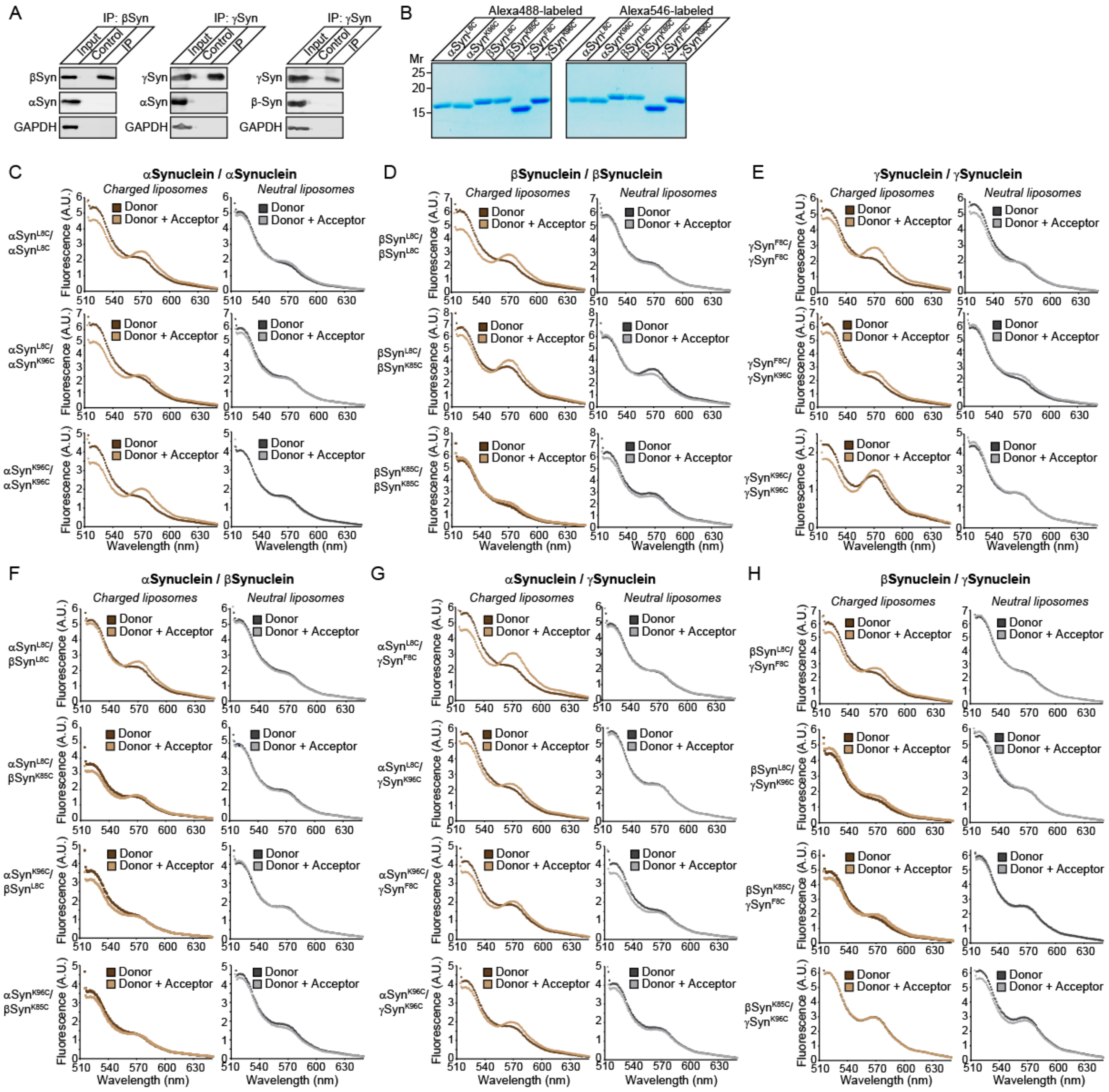


**Supplementary Figure S3. Synucleins interact with each other.**

(A) Synuclein interactions. Equal amounts of cDNA for the indicated synucleins were co-transfected into HEK293T cells. Synucleins in cell lysates were immunoprecipitated with preimmune serum (control) or with antibodies to βSyn or γSyn (IP). The immunoprecipitates were analyzed by immunoblotting with antibodies to αSyn, βSyn, or γSyn, and GAPDH as negative control.

(B) SDS/PAGE analysis of purified Alexa 488- or Alexa 546-labeled recombinant α-synuclein proteins (5 μg of protein per lane).

(C-H) Fluorescence resonance energy transfer (FRET) between donor and acceptor α-synuclein. Alexa 488-labeled donor αSyn (2.5 μg) was incubated for 2 h with Alexa 546-labeled acceptor or unlabeled αSyn (2.5 μg) in the presence of 100 μg of charged (red spectra; composition: 70% PC, 30% PS) or neutral small unilamellar vesicles (blue spectra; composition: 100% PC) of 30 nm.

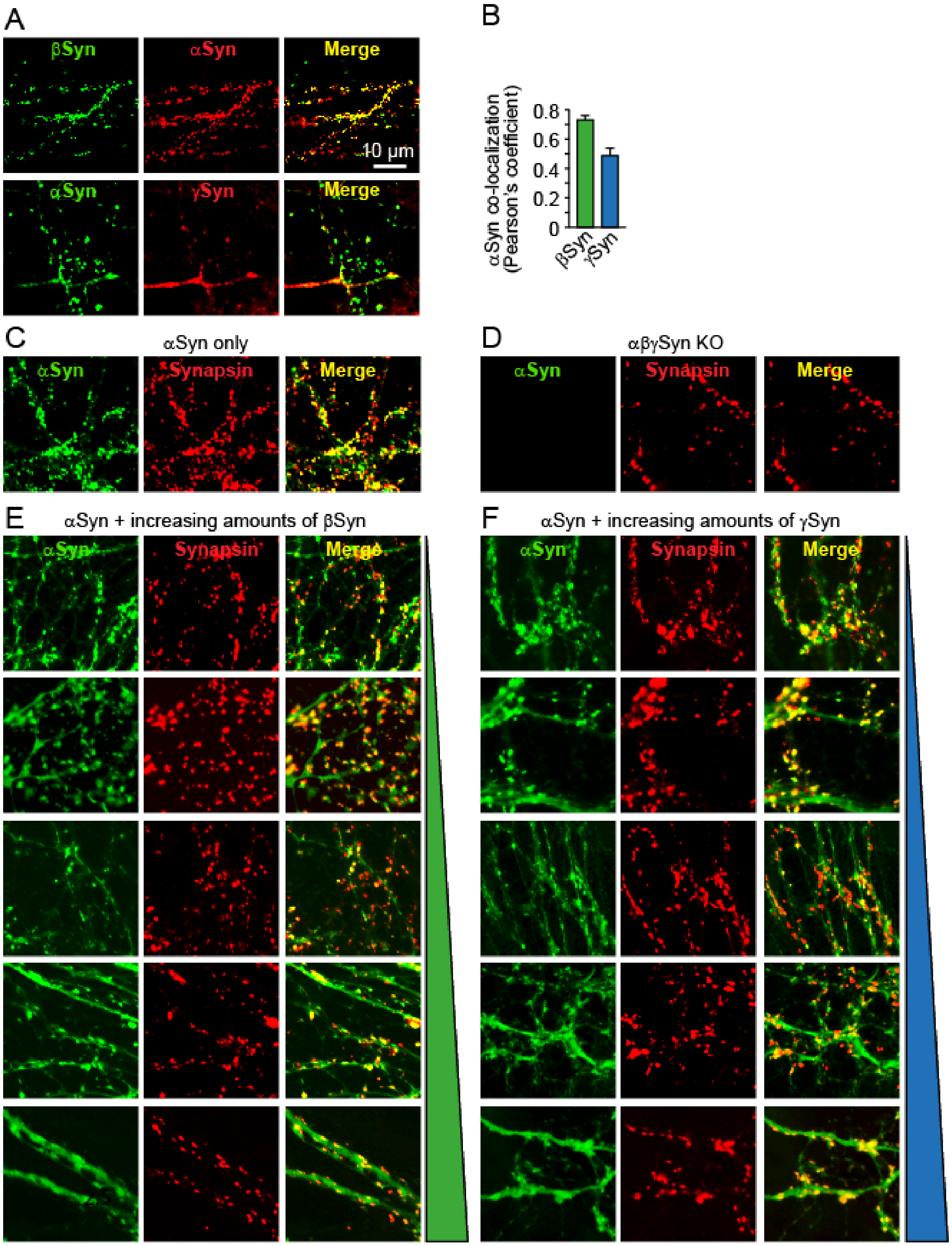


**Supplementary Figure S4. Decreased synaptic vesicle binding of αSyn in presence of βSyn or γSyn.**
(A, B) Primary cortical WT mouse neurons were analyzed at 27DIV for localization of αSyn, βSyn and γSyn. Colocalization was quantitated using Pearson's coefficient. Data are means ± SEM (n = 4 independent cultures).
(C-F) Primary cortical αβγ-synuclein knockout mouse neurons were infected with lentivirus expressing αSyn only (C), empty vector (D), or αSyn with increasing amounts of lentiviral vectors expressing βSyn (E) or γSyn (F). Neurons were analyzed at 21DIV for the indicated proteins. Colocalization was quantitated using Pearson's coefficient (see also Fig. 4A-D).

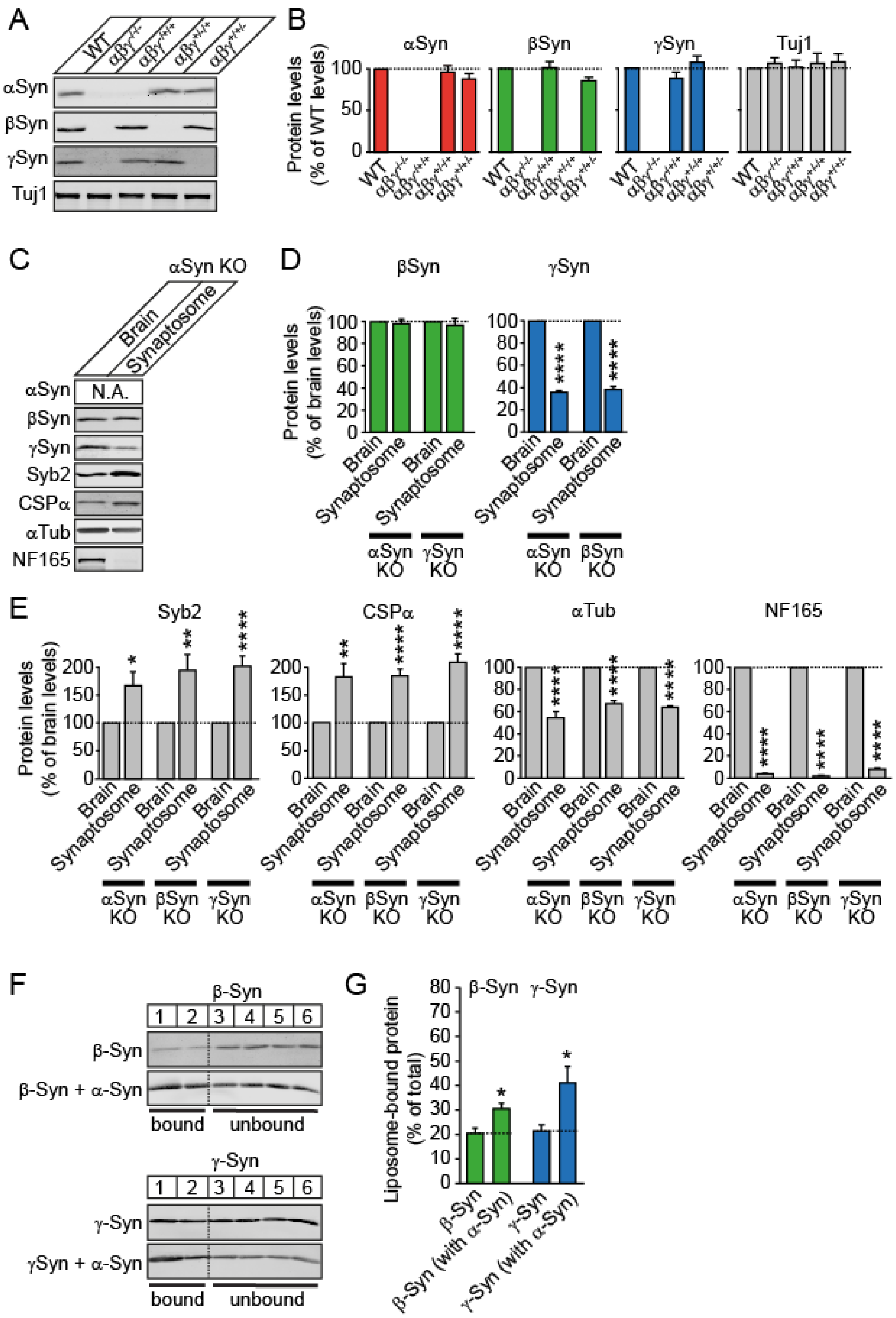


**Supplementary Figure S5. Decreased synaptic vesicle binding of αSyn in presence of βSyn or γSyn.**
(A, B) Total expression levels of synucleins in various genotypes**.** Brains of P40 WT mice were homogenized and 20 μg protein was separated by SDS-PAGE and analyzed by quantitative immunoblotting of the indicated protein. Data are means ± SEM (n = 6-8 mouse brains).
(C-E) Analysis of protein enrichment or depletion in synaptosomes. Synaptosomes were isolated from mouse brain homogenates of mice lacking αSyn, βSyn, or γSyn via subcellular fractionation. 20 μg protein of homogenate and synaptosomes were analyzed by quantitative immunoblotting to the indicated proteins (Syb2, synaptobrevin-2; αTub, α-tubulin; NF165, neurofilament of 165 kDa). Data are means ± SEM (* $p < 0.05$, ** $p < 0.01$, **** $p < 0.0001$ by Student t test; n = 6-8 mice; see also Figure 4E and 4F).
(F, G) Co-flotation of synucleins. Liposome binding of βSyn or γSyn was analyzed in absence or presence of equal amounts of αSyn by a flotation assay. Flotation of βSyn or γSyn with liposomes was quantified as the

sum of the top 2 fractions, plotted as the percentage of total synuclein in the gradient. Data are means ± SEM (* $p < 0.05$ by Student's t test; n = 6-15 independent experiments).
